# Development and Validation of a Method for Human Papillomavirus Genotyping Based on Molecular Beacon Probes

**DOI:** 10.1101/330936

**Authors:** Jiangfeng Lyu, Yuefeng Yu, Xuyi Ren

**Affiliations:** Research and Development Centre, Hangzhou DiAn Medical Laboratory, Hangzhou, Zhejiang, China

## Abstract

We describe a new assay system for the genotyping of human papillomavirus (HPV) based on linear-after-the-exponential-PCR and melting curve analysis. This system can detect and identify the 23 most common HPV strains (types 6, 11, 16, 18, 31, 33, 35, 39,42, 45, 51, 52, 53, 56, 58, 59, 66, 68, 70, 73, 81, 82, and 83) in two sealed reaction tubes within 2 h. The sensitivity and specificity of this new system was validated using cloned HPV DNA and clinical samples. The detection limit was 5–500 copies/reaction depending on the genotype, and no cross-reactivity was observed with any other low-risk HPV or pathogen that was commonly found in the female genital tract. When compared with the HPV GenoArray test kit, the testing of 1104 clinical samples produced a good overall agreement between the two methods of 98.37% (95% CI: 97.44%–98.97%) and a kappa value of 0.954. Thus, this new HPV genotyping assay system represents a simple, rapid, universally applicable, sensitive, and highly specific detection methodology that should be useful for cervical lesion screening and, thus, is potentially of great value in future clinical applications.

## Introduction

Cervical cancer is one of the most prevalent cancers among women worldwide, which has an estimated 527,624 new cases and 265,672 deaths in 2012, with over 85% occurring in developing countries (1). Cytology-based screening—the Papanicolaou (Pap) test—has helped in reducing the incidence of invasive cervical cancer over the past 50 years (2). However, the sensitivity of single Pap tests is relatively low, which leads to the missed detection of some cervical cancers and cervical intraepithelial neoplasia grades II (CIN II) and III (CIN III) (3). Cervical cancer is usually caused by persistent infection with human papillomavirus (HPV), particularly with high-risk genotypes including HPV16, 18, 31, 33, 35, 39, 45, 51, 52, 56, 58, 59, 66, and 68 (1, 4, 5). Thus, molecular assays have been developed for the detection of HPV DNA in cervical specimens and are already widely used in clinics. These molecular methods are more sensitive than the Pap test for the detection of CIN II and CIN III (6, 7). Currently, the most common detection and genotyping method for HPV in China is the HPV GenoArray test (HPV GenoArray test kit; Hybribio Ltd, Hong Kong). However, the main drawbacks of these assays are that they are prone to contamination and are time consuming (8). In addition, the diagnosis of borderline cases is difficult because of the read-outs being solely based on direct visualization (8).

On the basis of these principles, in this study, we developed a new test that uses linear-after-the-exponential (**L**ATE)-**P**CR and **M**elting curve analysis (LPM) for the DNA detection and genotyping of the 23 most common HPV strains (types 6, 11,16, 18, 31, 33,35, 39, 42, 45, 51, 52, 53, 56, 58, 59, 66, 68, 70, 73, 81, 82, and 83). We evaluated the analytical characteristics and clinical performance of the new test and found that it is faster and more convenient with more abundant operation than the current commercially available kits.

## Materials and Methods

### Sample collection

One liquid-based cytology (LBC) sample was collected from each of 1108 patients aged 19–89 years (median age, 43 years) who visited the DiAn Medical Laboratory for routine HPV screening or genotyping from August to December 2017. Additional 920 samples (40 of each HPV type including HPV types 6, 11, 16, 18, 31, 33, 35, 39, 42, 45, 51, 52,53, 56, 58, 59, 66, 68, 70, 73, 81, 82, and 83), which tested positive using the HPV GenoArray test kit or by type-specific PCR sequencing (for HPV types 70, 73, 82, and 83), were selected and stored at -80°C until needed. The selected HPV-positive samples were collected from 920 patients aged 23–92 years (median age, 45 years) from January 2016 to May 2017. All samples were collected using a cervical cytobrush and placed into tubes with 2 ml of PreservCyt solution (U.S. Cyt Company) along with the cytobrush. The collection of all cervical samples was approved by the ethics committee of DiAn Medical Laboratory.

### DNA extraction

The LBC samples were used for total genomic DNA extraction using a QIAamp DNA Mini kit (Qiagen, Hilden, Germany) according to the manufacturer’s protocol. DNA was eluted from the columns in 100 µL of AE buffer and then stored at -20°C.

### Primers and molecular beacon probes design

Various consensus primers have been reported for the amplification of a specific fragment of L1 the region of HPV DNA, including GP5+/6+, MGP, MY09/11, and PGMY09/11 (8-11). After considering amplification efficiency and the need for molecular beacon designation, GP6+/MY11 was chosen as the LATE-PCR primer set to generate the single strand DNA template for hybridization of the corresponding molecular beacon probe. Molecular beacon probes were designed and evaluated with the DNA folding program mfold (http://unafold.rna.albany.edu/?q=DINAMelt/Quickfold), and the probe–target hybrid folding program DINAMelt (http://unafold.rna.albany.edu/?q=DINAMelt/Two-state-melting) was used to predict the possible hybrid structures and T_m_ values. Sample quality control was evaluated by amplifying a 100-base pair (bp) fragment of the glyceraldehyde 3-phosphate dehydrogenase (GAPDH) gene using the primers G100f 5′-CTGCTCACATATTCTGGAGGAG-3′ (forward) and G100r 5′-AAAAGCAGCCCTGGTGACC-3′ (reverse). All molecular beacon probe sequences and corresponding targets are listed in Table 1.

**Table 1.**
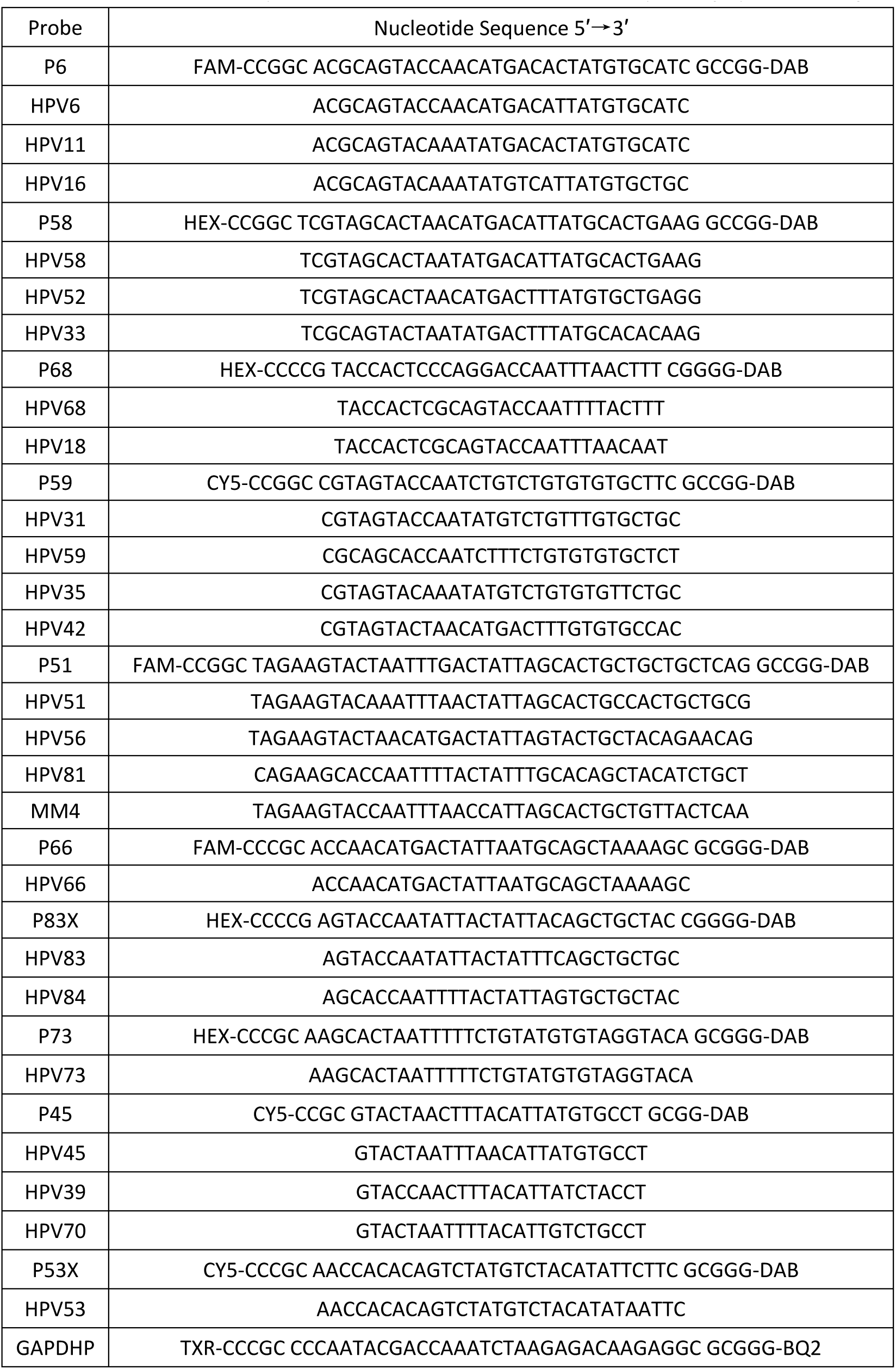
Molecular beacon probes used in LATE-PCR and their corresponding hybridized target sequences

### LATE-PCR

LATE-PCR was performed in a volume of 40 μl with 5 μl of extracted DNA as template. Every sample was tested with two reactions A and B. Each reaction comprised 1× PCR Buffer (Takara, Japan), 2 mM MgCl_2_ (Promega, USA), 12.5 nM dU plus deoxynucleotide triphosphates (dNTPs, Takara), 37.5 nM limiting primer MY11 (5′-GCMCAGGGWCATAAYAATGG-3′), 500 nM excess primer GP6+ (5′-GAAAAATAAACTGTAAATCATATTC-3′), two units of HS Taq polymerase (Takara), and one unit of Uracil DNA Glycosylase (UNG, Takara). Probes for reaction A were 25 nM of probe P6, 15 nM of probes P58 and P68, and 20 nM of probe P59, and those for reaction B were 25 nM of probes P51 and P66, 15 nM of probes P73 and P83, and 20 nM of probes P45 and P53. Reaction B also contained internal control primers and probe, those being 50 nM of forward primer G100f, 50 nM of reverse primer G100r, and 20nM of probe GAPDHP. All primers and probes were synthesized and purified by Invitrogen Life Technologies (Shanghai, China). The thermal profile conditions used on a LightCycler480 (Roche, Germany) were as follows: 50°C for 10 min and 95°C for 5 min, followed by 95°C for 15 s, 45°C for 30 s, and 72°C for 30 s for 55 cycles. This was followed by a melt starting at 37°C with 1°C increments at 30 s intervals to 85°C. Fluorescence signals were collected and analyzed in four different channels: FAM (465-510), HEX (533-580), TXR (533-610), and CY5 (618-660). To generate the corresponding T_m_ values, melt-curve analysis using the first derivative of fluorescence at temperatures between 37°C and 85°C was used to process the fluorescence data of the probe–target hybridizations.

### Determination of reference value

The 920 HPV-positive samples were further tested using the LPM genotyping assay. Data were collected as T_m_ values. The nonparametric method was used to calculate the reference interval. Upper and lower limit values of the reference interval were calculated using the nonparametric method (percentile estimation method), and points corresponding to 95% of the distribution were sought. Results were evaluated using SPSS 16.0. The Kolmogorov–Smirnov Z-test was used as a normal distribution test.

### Analytical sensitivity and specificity

To assess analytical sensitivity, recombinant plasmids containing the cloned PCR products of the full-length L1 gene were generated using appropriate primers for each HPV type from DNA extracted from the positive samples screened with the HPV GenoArray test. The PCR products were purified and inserted into a pMD19-T vector (Takara) following the manufacturer’s instructions. Cloned PCR products were then sequenced on an ABI3130 genetic analyzer (Applied Biosystems, USA). Sequences were subjected to BLAST search in the GenBank database and were verified as being 100% homologous with the target sequences. Plasmid concentrations were quantified with a NanoDrop1000 Spectrophotometer (NanoDrop, DE). Copy numbers of the cloned HPV L1 genes were derived from the molecular weights of the recombinant plasmids and samples diluted with TE buffer (pH 8.0) to generate concentration gradient ranging from 1 to 10^6^ copies/ml. Each dilution also contained the same amount (40 ng/ml) of human genomic DNA. Five μl of each plasmid dilution were used as templates in LATE-PCR and each HPV type run in 20 replicate reactions per dilution.

To evaluate specificity, DNA was purified from LBC samples that were positive for *Neisseria gonorrhoeae* (ATCC49226), *Candida albicans* (ATCC90028), *Escherichia Coli* (ATTCC25922), *Staphylococcus aureus* (ATTCC25923), Herpes simplex virus type II, *Treponema pallidum, Ureaplasma urealyticum,* and *Mycoplasma hominis* using the QIAamp DNA Mini kit. Five microliter of HPV26, 40, 43, 44, 54, 61, 67, 69, 71, and 72 plasmids at a concentration of 10^8^ copies/ml and purchased from National Institutes for Food and Drug Control were used as templates in LATE-PCR.

### Comparison of the HPV GenoArray test with the LPM genotyping assay

Both assays were used to independently test 1108 LBC samples in order to compare HPV genotyping results. The GenoArray test is an L1 consensus primer-based PCR assay and is capable of amplifying 21 HPV genotypes, including types 16, 18, 31, 33, 35, 39, 45, 51, 52, 56, 58, 59, 68, 53, 66, 6, 11, 42, 43, 44, and 81. This includes two HPV genotypes that were out of the detection range of the LPM assay (i.e., low-risk HPV genotypes 43 and 44) and four genotypes that could be detected, but not quantified, using the LPM assay (genotypes 70, 73, 82, and 83). All GenoArray detection procedures were performed according to the manufacturer’s instructions, and LPM genotyping was conducted as described above.

### Type-specific sequencing analysis

Sequencing was performed when there was a discrepancy in the results between the two assays as described by Peng *et al.* (3). Briefly, the PGMY09/11 primer set was used to amplify a specific fragment of the L1 gene from purified DNA. HPV genotype was assigned by sequencing of the amplified fragments using a type-specific sequencing primer. Sequencing was performed using the BigDye Terminator v3.1 cycle sequencing kit with AmpliTaq DNA polymerase (Applied Biosystems) according to the protocols provided by the manufacturer. The sequences obtained were compared with known HPV sequences using the BLAST tool (http://www.ncbi.nlm.nih.gov/BLAST).

## Results

### Reference T_m_ values of each HPV type

Forty HPV-positive patient samples for each HPV type were used to establish a statistically significant reference interval. All samples were tested with the LPM assay as described above to produce 40 T_m_ values for each HPV type. Each T_m_ value distribution was assessed as to whether it exhibited a Normal distribution and then reference Tm values were generated and evaluated using SPSS 16.0. Reference T_m_ values for each HPV type are shown in Table 2. In separate reactions (A or B), there were no overlapping reference T_m_ intervals between the different HPV types within a single fluorescence channel.

**Table 2.** T_m_ value of each HPV type

| Reaction | Fluorescent labeling | Excitation and Emission wavelengths (nm) | HPV type | T<sub>m</sub> Value |
| --- | --- | --- | --- | --- |
|  |  |  | 45 | 49.82–51.00 |
| A | FAM | 465-510 | 16 | 53.33–55.45 |
|  |  |  | 11 | 58.00–59.41 |
|  |  |  | 6 | 64.44–67.22 |
|  | HEX | 533-580 | 68 | 47.09–48.63 |
|  |  |  | 18 | 51.40–53.00 |
|  |  |  | 33 | 54.69–55.90 |
|  |  |  | 52 | 57.11–59.13 |
|  |  |  | 58 | 61.77–63.89 |
|  | CY5 | 618-660 | 42 | 47.49–49.24 |
|  |  |  | 59 | 52.17–54.13 |
|  |  |  | 35 | 55.62–56.59 |
|  |  |  | 31 | 59.93–64.00 |
| B | FAM | 465-510 | 81 | 50.65–52.10 |
|  |  |  | 56 | 53.84–55.35 |
|  |  |  | 82 | 57.03–58.18 |
|  |  |  | 51 | 59.66–61.20 |
|  |  |  | 66 | 64.98–66.49 |
|  | HEX | 533-580 | 83 | 57.79–59.75 |
|  |  |  | 73 | 63.72–65.41 |
|  | CY5 | 618-660 | 39 | 45.35–47.35 |
|  |  |  | 70 | 54.89–56.04 |
|  |  |  | 53 | 60.46–66.92 |

### Analytical sensitivity and specificity of the LPM assay

LPM assay sensitivity was assessed with serial dilutions of plasmids harboring the L1 gene and containing 1 to 10^6^ copies/ml and the same amount (40 ng/ml) of human genomic DNA. The analytical sensitivity or limit of detection of the assay, defined as the lowest concentration (target gene copy number per reaction) where ≥ 95% of test runs produced positive results, was estimated using these serial dilutions. The limit of detection for HPV6, 11,16, 18, 33, 35, 45, 52, 56, 58, 66, 68, 73, and 81 was 50 copies/reaction; for HPV31, 42, 51, 53, and 59 was 500 copies/reaction; and for HPV39, 70, 82, and 83 was 5 copies/reaction.

### HPV genotyping results of GenoArray test and LPM assay

These two assays can detect largely overlapping sets of HPV genotypes, and HPV genotyping results were compared using 1108 LBC samples; the results were mostly concordant or compatible. Among the 1108 samples tested, four generated no melting peak in the control channel of the LPM assay, meaning that there was either insufficient DNA to be amplified or a PCR inhibitor was present in the extracted DNA. Therefore, comparable results were obtained for a total of 1104 samples. With respect to the overall detection of HPV DNA, the absolute agreement between the tests was 98.37% (95% CI: 97.44%–98.97%) and the kappa value of 0.954, indicating good agreement. The positive concordance rate was 98.94% (95% CI: 98.00%–99.44%) and the negative concordance rate was 96.43% (95% CI: 93.35–98.11%; Table 3). Subsequently, a comparison of the specific identification of individual HPV genotypes was made and the results are summarized in Table 4. The two assays were in perfect agreement for almost all HPV types within the scope of test, with kappa values of >0.75 except for HPV45, for which the two assays exhibited only moderate agreement (kappa values, 0.40–0.75). The LPM assay detected HPV 45 in significantly more samples than the GenoArray, indicating that the LPM assay was more sensitive in this respect.

**Table 3.** Comparison of results for the overall detection of HPV DNA by the two assays

| LPM assay result | GenoArray assay result |  |  |
| --- | --- | --- | --- |
|  | Positive | Negative | Total |
| Positive | 843 | 9 | 852 |
| Negative | 9 | 243 | 252 |
| Total | 852 | 252 | 1104 |

**Table 4.** Interassay agreements for individual HPV genotypes detected by two assays

| HPV<br>genotype | No. of samples positive by |  |  | Absolute<br>agreement (%) | Kappa<br>value |
| --- | --- | --- | --- | --- | --- |
|  | LPM | GenoArray/Sequencing | Both |  |  |
| 6 | 41 | 41 | 40 | 99.82 | 0.975 |
| 11 | 25 | 26 | 24 | 99.73 | 0.940 |
| 16 | 141 | 144 | 140 | 99.55 | 0.980 |
| 18 | 42 | 43 | 36 | 98.82 | 0.841 |
| 31 | 24 | 26 | 24 | 99.82 | 0.959 |
| 33 | 29 | 37 | 29 | 99.28 | 0.875 |
| 35 | 25 | 26 | 25 | 99.91 | 0.980 |
| 39 | 54 | 55 | 53 | 99.73 | 0.971 |
| 42 | 20 | 25 | 19 | 99.37 | 0.841 |
| 45 | 23 | 9 | 9 | 98.73 | 0.557 |
| 51 | 51 | 53 | 50 | 99.64 | 0.960 |
| 52 | 161 | 158 | 156 | 99.37 | 0.974 |
| 53 | 81 | 81 | 80 | 99.82 | 0.987 |
| 56 | 60 | 61 | 59 | 99.73 | 0.974 |
| 58 | 103 | 101 | 100 | 99.64 | 0.978 |
| 59 | 34 | 41 | 33 | 99.18 | 0.876 |
| 66 | 39 | 38 | 38 | 99.91 | 0.987 |
| 68 | 14 | 17 | 14 | 99.73 | 0.902 |
| 70 | 15 | 14 | 14 | 99.91 | 0.965 |
| 73 | 9 | 8 | 8 | 99.91 | 0.941 |
| 81 | 60 | 60 | 59 | 99.82 | 0.982 |
| 82 | 22 | 20 | 20 | 99.82 | 0.951 |
| 83 | 11 | 10 | 10 | 99.91 | 0.952 |

Among the 1104 cases compared, 73 exhibited discordant GenoArray and LPM assay results and six exhibited discordant sequencing and LPM assay for HPV70, 73, 82, and 83. The 73 discordant cases were further analyzed by type-specific PCR followed by sequencing (Table 5). After sequencing, 40 cases matched the LPM assay result and 31 matched by GenoArray, whereas in two cases, the sequencing disagreed with both assays.

**Table 5.** Overview of the discrepancy analysis of discordant genotypes reassessed by sequencing

| HPV type | No. of samples with discordant types |  |  |  |  |
| --- | --- | --- | --- | --- | --- |
|  | Total | Detected by<br>GenoArray | Confirmed by<br>sequencing | Detected by<br>LPM assay | Confirmed by<br>sequencing |
| 6 | 2 | 1 | 0 | 1 | 1 |
| 11 | 3 | 2 | 1 | 1 | 1 |
| 16 | 5 | 4 | 2 | 1 | 1 |
| 18 | 13 | 7 | 5 | 6 | 6 |
| 31 | 2 | 2 | 2 | 0 | 0 |
| 33 | 8 | 8 | 6 | 0 | 0 |
| 35 | 1 | 1 | 1 | 0 | 0 |
| 39 | 3 | 2 | 2 | 1 | 1 |
| 42 | 7 | 6 | 1 | 1 | 1 |
| 45 | 14 | 0 | 0 | 14 | 14 |
| 51 | 4 | 3 | 1 | 1 | 1 |
| 52 | 7 | 2 | 0 | 5 | 5 |
| 53 | 2 | 1 | 1 | 1 | 1 |
| 56 | 3 | 2 | 2 | 1 | 1 |
| 58 | 4 | 1 | 1 | 3 | 3 |
| 59 | 9 | 8 | 7 | 1 | 1 |
| 66 | 1 | 0 | 0 | 1 | 1 |
| 68 | 3 | 3 | 1 | 0 | 0 |
| 81 | 2 | 1 | 0 | 1 | 1 |

## Discussion

Since the discovery of a relationship between HPV and cervical cancer, various methods have been applied to HPV detection and genotyping, all of which mainly rely on molecular biology techniques and have improved the diagnosis and management of early cervical abnormalities and precancerous lesions. The Hybrid Capture 2 system (HC2, Digene Corp., USA) is based on signal amplification and detects 13 HR-HPV types (16, 18, 31, 33, 35, 39, 45, 51, 52, 56, 58, 59, and 68) and five LR-HPV types (6, 11, 42, 43, and 44). While it can distinguish high-risk from low-risk types, it is incapable of HPV genotyping (9). However, identifying HPV types is clinically important. International consensus is that persistent infection of HR genotypes, such as HPV16, 18, 31, 33, 35, 39, 45, 51, 52, 56, 58, 59, and 66, can lead to cervical cancer, whereas infections with LR viruses cause only benign cervical tissue alterations or genital warts (10). If specific HPV genotypes could be confirmed, then a more efficient management can be implemented on the basis of the HPV type identified, and novel strategies based on the use of HPV DNA assays for primary cervical screening are increasingly being recommended (11). In addition, promising new broad-spectrum HPV vaccines are in development (12). Bivalent (against types 16 and 18), quadrivalent (against types 6, 11, 16, and 18) and nonavalent (against types 6, 11, 16, 18, 31, 33, 45, 52, and 58) vaccines have been developed and proven safe and efficiently reduced the incidences of HPV infections, genital warts, and HPV-attributed precancerous lesions (13-19). Thoughtful integration of vaccination and screening will reduce not only cervical cancer risk but also over-treatment for abnormalities destined to resolve.

Currently, the most commercially used method for HPV genotyping in China is the HPV GenoArray test. This method is based on reverse-line blot technology, in which PCR products are hybridized to HPV type-specific probes on a membrane. It is an L1 consensus primer-based PCR assay and is capable of amplifying 21 HPV genotypes, including 13 HR types (16, 18, 31, 33, 35, 39, 45, 51, 52, 56, 58, 59, and 68), two probable HR types (types 53 and 66) and six LR and unknown-risk types [types 6, 11, 42, 43, 44, and CP8304 (HPV-81)]. Numerous studies have shown that the GenoArray test is highly reproducible and exhibits excellent clinical sensitivity for these HPV types; therefore, it was considered to be suitable for comparison with the LPM assay in this study (8, 20). However, the main drawbacks of GenoArray are its vulnerability to contamination and its time-consuming nature (8). In addition, it is sometimes difficult to give an unambiguous diagnosis for borderline cases because read-out is based only on direct visualization.

Real-time PCR based on Taqman probe has been used for pathogen detection and identification in clinical samples with several targets because of the limited fluorophores (21-24). However, it is often desirable to detect and identify multiple types of pathogen that might be present in a clinical sample. Molecular beacons labeled with different fluorophores and specific to different target sequences can be combined to simultaneously detect and identify multiple targets in a single assay tube (25). Chakravorty *et al.* were able to achieve rapid universal identification of bacterial pathogens from a list of 111 species in 64 different genera using a molecular beacon melting temperature signature technique (26). Additionally, El-Hajj *et al.* described a molecular beacon system that can successively distinguish 27 different species of Mycobacteria (27). These assays identify multiple targets not only by distinguishing between the fluorescence wavelengths of each different molecular beacon but also by accounting for differences in the melting temperatures of the probe sequence hybridized to their targets. In this study, a new HPV detection and genotyping system—the LPM assay—was developed based on molecular beacons and its performance on clinical samples was validated by comparing with the GenoArray test. An excellent agreement (kappa = 0.954; 95% CI, 97.44%–98.97%) between the LPM assay and GenoArray was obtained for HPV genotyping. Compared with GenoArray, the sensitivity, specificity and accuracy of the LPM assay were 98.94 %, 96.43 % and 98.37 %, respectively. This indicates HPV genotypes can be successfully identified by the LPM assay with high accuracy. As to individual HPV genotypes, the GenoArray and LPM assays were in near-perfect agreement for the detection of almost all HPV types within the scope of test, with kappa values of >0.75, except for HPV45, where the two assays exhibited moderate agreement (kappa values, 0.40–0.75). The LPM assay detected HPV type 45 in significantly more samples than the GenoArray, indicted the LPM assay was more sensitive for the detection of this genotype. Of the 1104 LBC samples tested with the LPM assay, only 33 cases were a mismatch with GenoArray or type-specific sequencing. These results indicate that the LPM method can successfully detect HPV infections and identify HPV genotypes with a high degree of accuracy. Additional practical advantages with the LPM assay are that it costs less (less than 1.5 US dollars per sample) and is faster (<2 h) than the GenoArray assay and that because LATE-PCR and fluorescence measurement is conducted in sealed tubes, carryover contamination cannot occur.

In conclusion, the LPM assay exhibited similar clinical performance for genotyping as the HPV GenoArray test and seemed to be a promising alternative approach for HPV testing due to its high level of automation. This new assay could be a useful tool for both primary screening for cervical cancer and for the triage of women exhibiting abnormal cytological results. It offers high sensitive, cost-effectiveness high throughput that could facilitate the rapid and specific detection of 23 high- and low-risk HPV genotypes.

## Acknowledgements

This work was supported by DiAn Medical Laboratory. The funders had no role in the study design, data collection, interpretation, or the decision to submit the work for publication.

We would like to thank Lei Zhou for the English language review. The authors declare that there are no conflicts of interest.

